# Skeletal Muscle Transcriptome Alterations Related to Physical Function Decline in Older Mice

**DOI:** 10.1101/2021.05.17.444371

**Authors:** Ted G. Graber, Rosario Maroto, Jill Thompson, Steve Widen, Zhaohui Man, Megan L. Pajski, Blake B. Rasmussen

**Affiliations:** Dept. of Physical Therapy, East Carolina University; Dept. of Nutrition and Metabolism, University of Texas Medical Branch; Next Generation Sequencing Core Facility, University of Texas Medical Branch; Bioinformatics and Analytics Research Collaborative, University of North Carolina-Chapel Hill

**Keywords:** muscle, transcriptome, aging, mice, functional performance, neuromuscular

## Abstract

One inevitable consequence of aging is the gradual deterioration of physical function and exercise capacity, driven in part by the adverse effect of age on muscle tissue. Our primary purpose was to determine the relationship between patterns of gene expression in skeletal muscle and this loss of physical function. We hypothesized that some genes changing expression with age would correlate with functional decline, or conversely with preservation of function. Male C57BL/6 mice (6-months old, 6m, 24-months, 24m, and 28+-months, 28m; all n=8) were tested for physical ability using a **<u>c</u>**omprehensive **f**unctional **<u>a</u>**ssessment **<u>b</u>**attery (CFAB). CFAB is a composite scoring system comprised of five functional tests: rotarod (overall motor function), grip strength (fore-limb strength), inverted cling (4-limb strength/endurance), voluntary wheel running (activity rate/volitional exercise), and treadmill (endurance). We then extracted total RNA from the tibialis anterior muscle, analyzed with Next Generation Sequencing RNAseq to determine differential gene expression during aging, and compared these changes to physical function. Aging resulted in gene expression differences >│1.0│ log2 fold change (multiple comparison adjusted p<0.05) in **219** genes in the 24m and in **6587** genes in the 28m. Linear regression with CFAB determined **253** differentially expressed genes strongly associated (R>0.70) with functional status in the 28m, and **22** genes in the 24m. We conclude that specific age-related transcriptomic changes are associated with declines in physical function, providing mechanistic clues. Future work will establish the underlying cellular mechanisms and the physiological relevance of these genes to age-related loss of physical function.

**Graphical Abstract:** 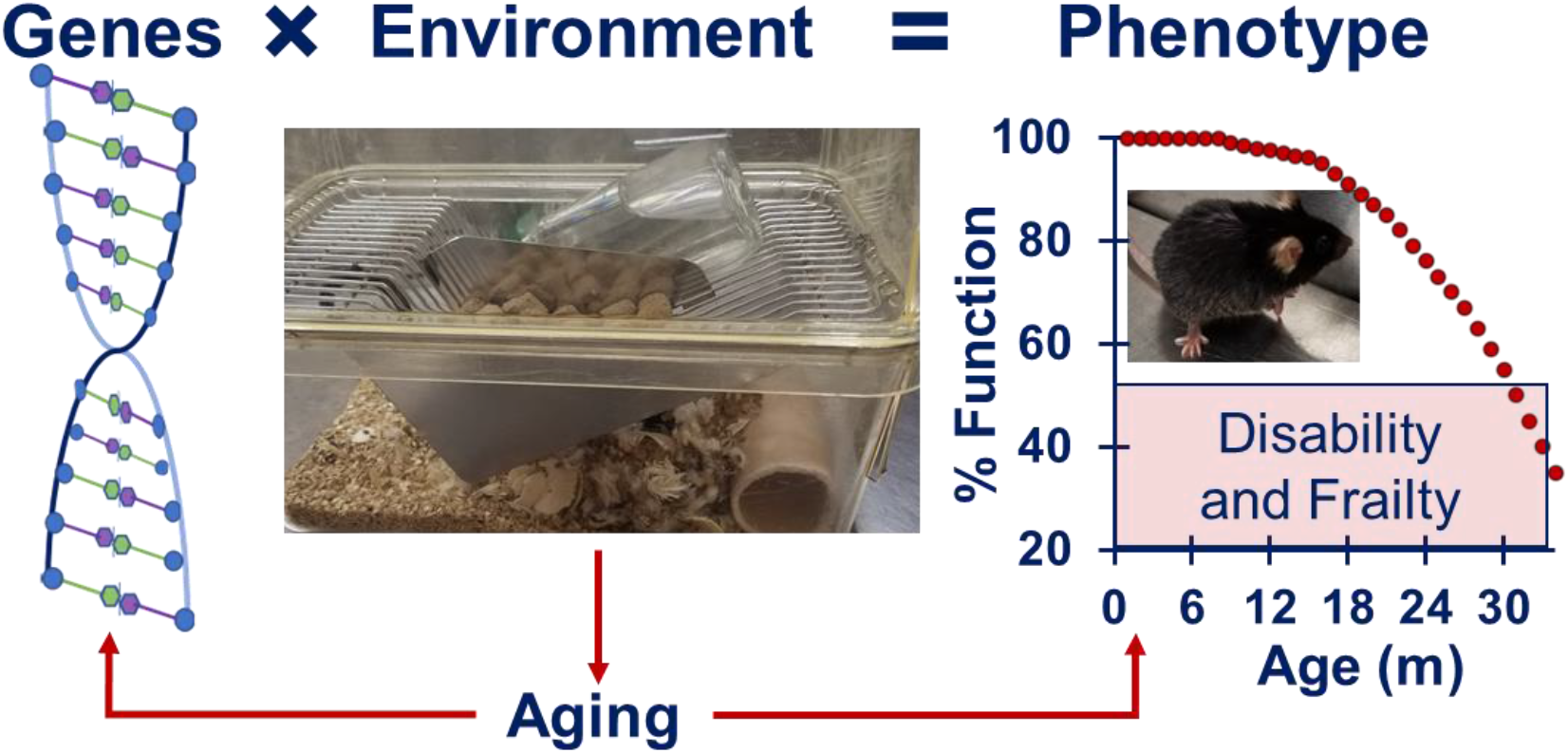

RNA sequencing of skeletal muscle from young and old mice were compared to physical function status obtained by performing a comprehensive functional assessment battery of tests. Between adulthood (6-months) and older age (28-months), 6707 genes were differentially expressed with 253 of these genes being significantly associated with physical function. Specific age-related changes to the skeletal muscle transcriptome are associated with a decline in physical function.

## Introduction

Aging results in the onset of the decline of physical function accompanied or predicated by loss of skeletal muscle mass and strength (sarcopenia). This loss of physical function and muscle health leads to reduced ability to perform activities of daily living, a lower quality of life, development of disability, eventual loss of independence, and increased mortality (Aversa, 2019; Tsekoura 2017; Billet, 2020). Sarcopenia and frailty (inability to maintain homeostasis) are linked in that most frail individuals are also sarcopenic. While we still do not know the exact etiology of sarcopenia, we do know that it is likely a multifactorial disease with a host of potential causes such as: disuse atrophy, neuromotor deficits including denervation, reduction in muscle quality (fat and fibrotic intrusion), alterations to key proteins and cell signaling, mitochondrial deficits, and many others (Pratt, 2020; Deschenes, 2011; Thompson, 2009; Coen, 2019, Narici, 2010). Understanding the molecular mechanisms leading to the age-associated skeletal muscle function will enable us to develop mitigation strategies for functional decline. In this study, our primary goal was to determine how changes in muscle gene expression during aging are related to physical function and exercise capacity. Our long-term goal is to utilize this novel data set to design experiments focused on identifying the underlying cellular mechanisms of sarcopenia.

To accomplish our goal, we used our Comprehensive Functional Assessment Battery (CFAB), a composite scoring system comprised of five different tests (rotarod, grip test, inverted cling, voluntary wheel running, treadmill), to measure physical function in 6-month-old (6m), 24-month-old (24m), and 28+month-old (28m) C57BL/6 mice (Graber, 2020). We then used Next Generation Sequencing (NGS) RNAseq to determine gene expression in the tibialis anterior (TA) muscles of these mice. By using linear regression of genes that changed expression in aging with physical function we were able determine the associations and note numerous genes in muscle that may play a critical role in declining physical ability.

We found thousands of genes with differential expression between 6m and 28m of age, versus only a couple hundred between the 6m and the 24m. Likewise, there were hundreds of genes changing expression with age that were strongly associated with CFAB in 28m, but far fewer in 24m. This discrepancy highlights potential acceleration of biological aging over those four months, that is also manifested in many indicators of physical function and muscle health (Graber 2015; Graber, 2020). GOrilla (Gene Ontology enRIchment anaLysis and visuaLizAtion tool) and GSEA (Gene Set Enrichment Analysis) were used to determine a number of highly enriched gene ontologies, which included cation transporters, and calcium transporters in particular (Eden, 2009; Subramanian, 2005). Overall, this novel data set establishes an initial framework for understanding how aging alters skeletal muscle gene expression and identification of specific muscle genes linked to the gradual, inevitable, progressive loss of physical function associated with sarcopenia.

## Results

### CFAB

The mice from this study demonstrated overall declining physical function with age, as measured with the CFAB component tests of rotarod, grip strength meter, inverted cling, treadmill, and voluntary wheel running (see **Figure 1**). The CFAB score was significantly different between groups (p<0.05). The mice in this study were randomly chosen from the larger cohort in our previously published work, refer to that work for a more complete discussion of the methods and results of the functional testing across the three age groups (Graber, 2020).

**Figure 1.**
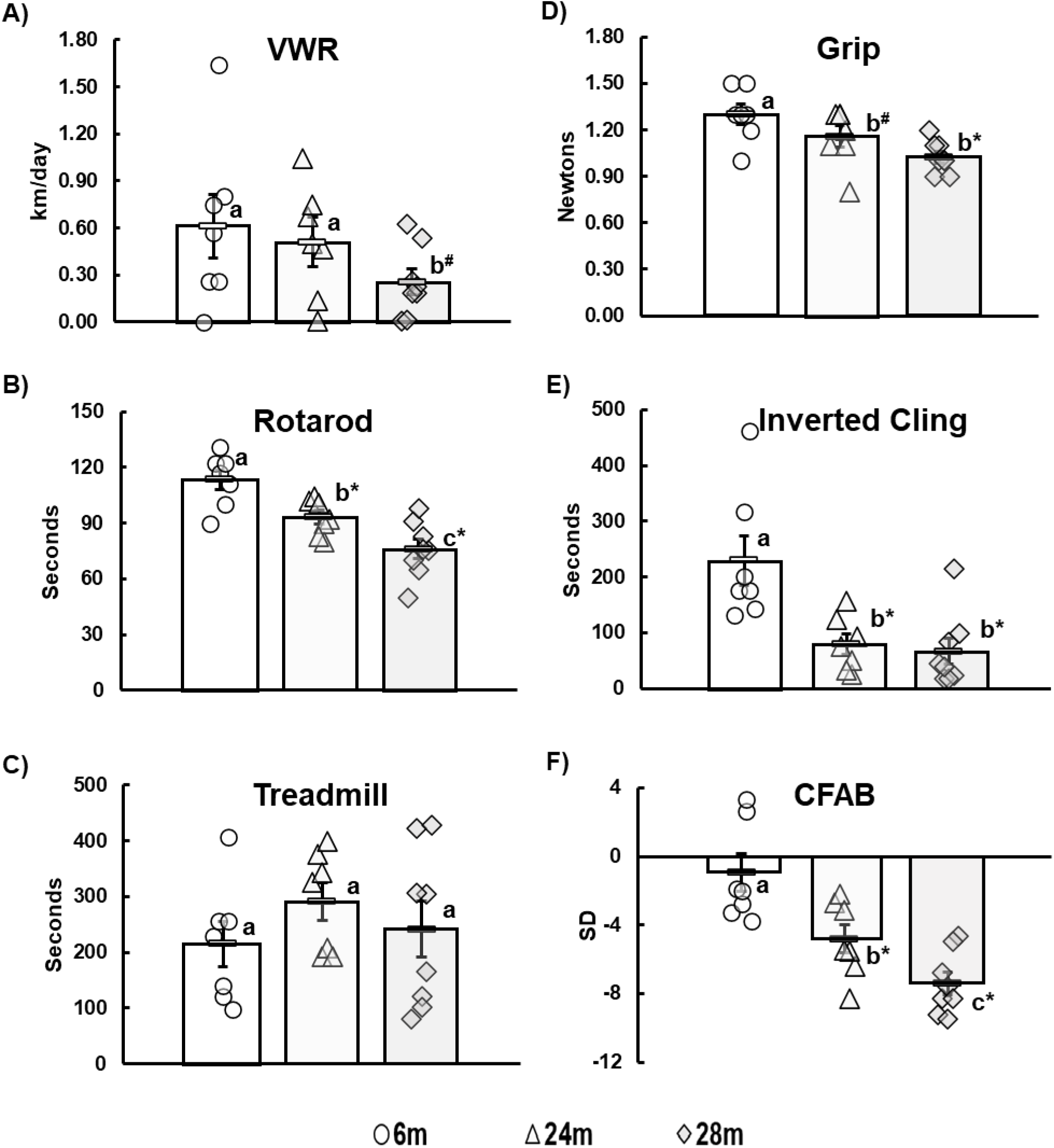
Declining Physical Function with Age. m=months, different letters indicate statistical significance, **\*** = p<0.05, ^**#**^ = 0.05<p<0.10. Each symbol indicates data from an individual mouse (circles 6m, n=7; triangles 24m, n=7; diamonds 28m, n=8). Statistics are from ANCOVA adjusted for body mass with Least Significant Differences post hoc testing.

### NGS RNAseq: (See the full raw dataset on GEO at GSE152133)

#### 28-month-old mice compared to 6-month-old mice

We determined that, overall, 6587 genes significantly changed (adj. p<0.05) with age in the 28m versus the 6m. By expanding to include genes changing with p≤0.05, there were 6707 genes, with 3153 downregulated (614, log2fc≤-1; adj. p≤0.05) and 3554 upregulated (615 with log2fc≥1; adj. p≤0.05). The top 50 gene expression changes between 6m and 28m are shown in a heatmap in **Figure 2** and a volcano plot showing separation of the gene sets in **Figure 3**. In **Table 1** we list the top 20 genes upregulated with age, and in **Table 2** the top 20 downregulated Genes, see **Table S1a** for all genes log2fc≥│1│and adj. p≤0.05. In **Figure 4A**, the 2D principal component analysis (PCA) scores plot indicates a separation between 28m and 6m clusters, with no overlap. This result was confirmed by using the supervised multivariate analysis based on a partial least squares-discriminate analysis (PLS-DA) (component 1, 6m, was 14% and component 2, 28m, was 56%).

**Table 1.**
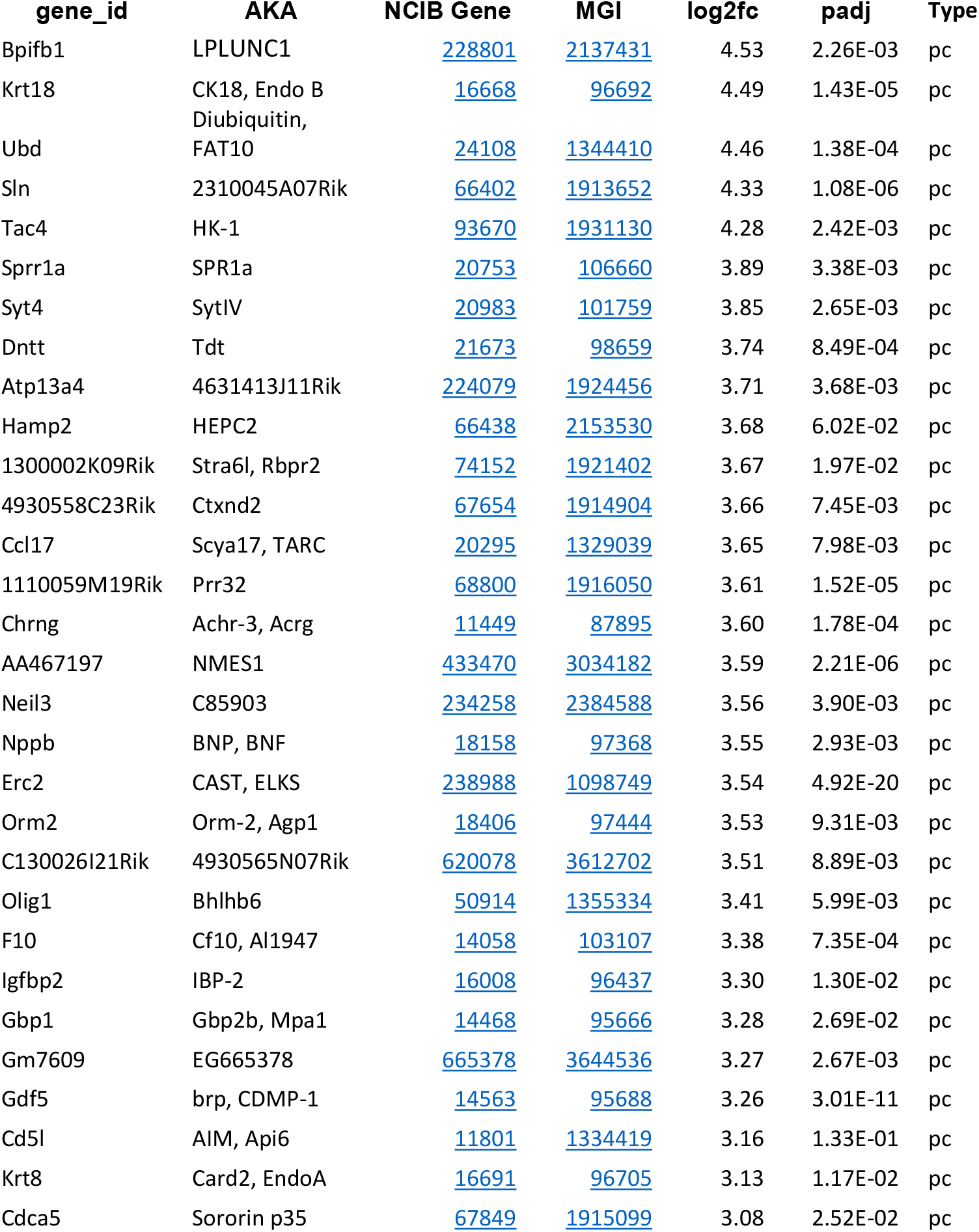
Top 30 Upregulated Aging Genes: 6m vs. 28m. AKA = also known as, NCIB Gene is from https://www.ncbi.nlm.nih.gov/gene, MGI = Mouse Genome Informatics from http://www.informatics.jax.org/marker, log2fc = log base 2 fold change, adj. p = multiple comparison adjusted p-value

**Table 2.**
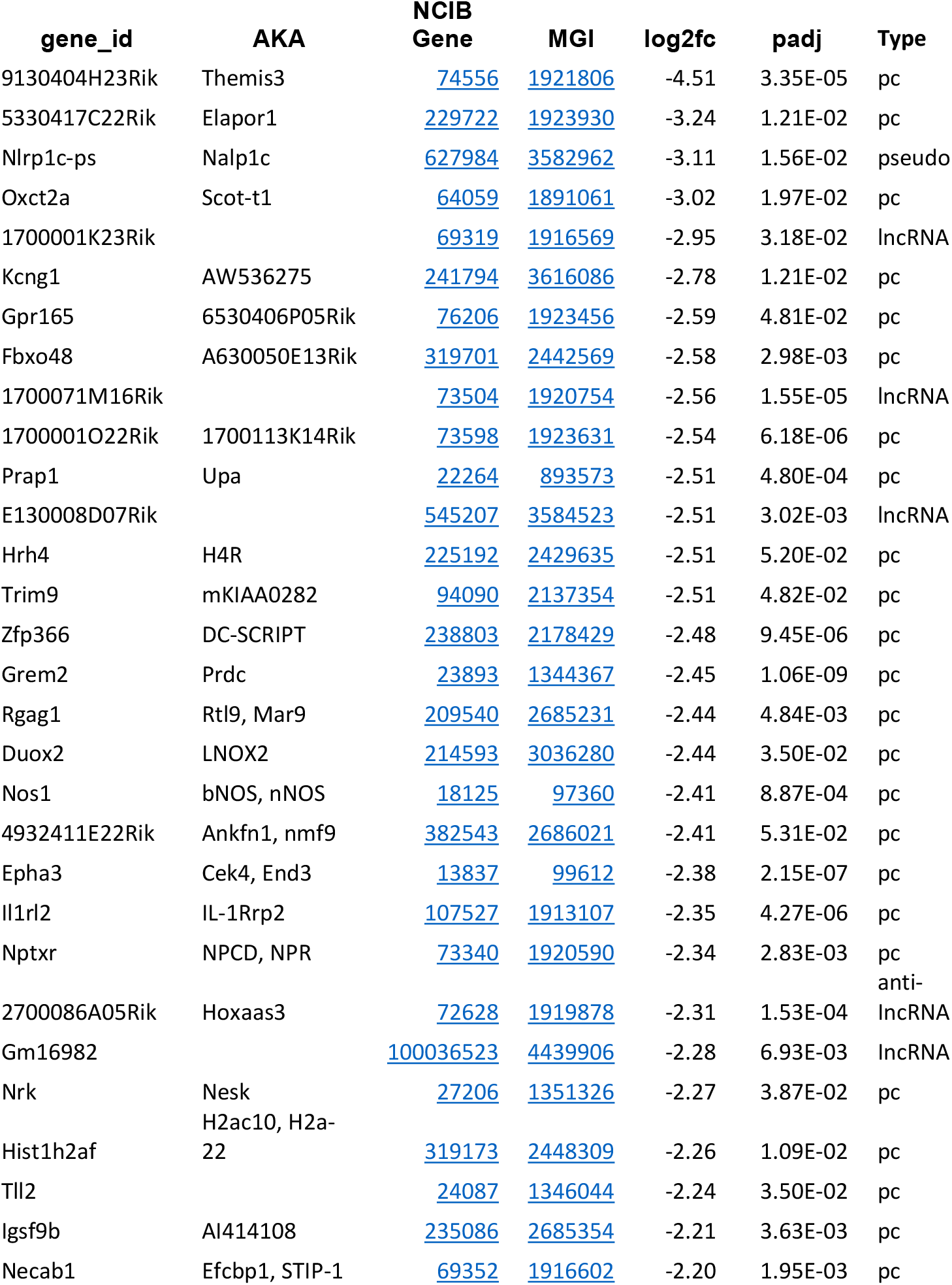
Top 30 Downregulated Aging Genes: 6m vs. 28m. AKA = also known as, NCIB Gene is from https://www.ncbi.nlm.nih.gov/gene, MGI = Mouse Genome Informatics from http://www.informatics.jax.org/marker, log2fc = log base 2 fold change, adj. p = multiple comparison adjusted p-value

**Figure 2.**
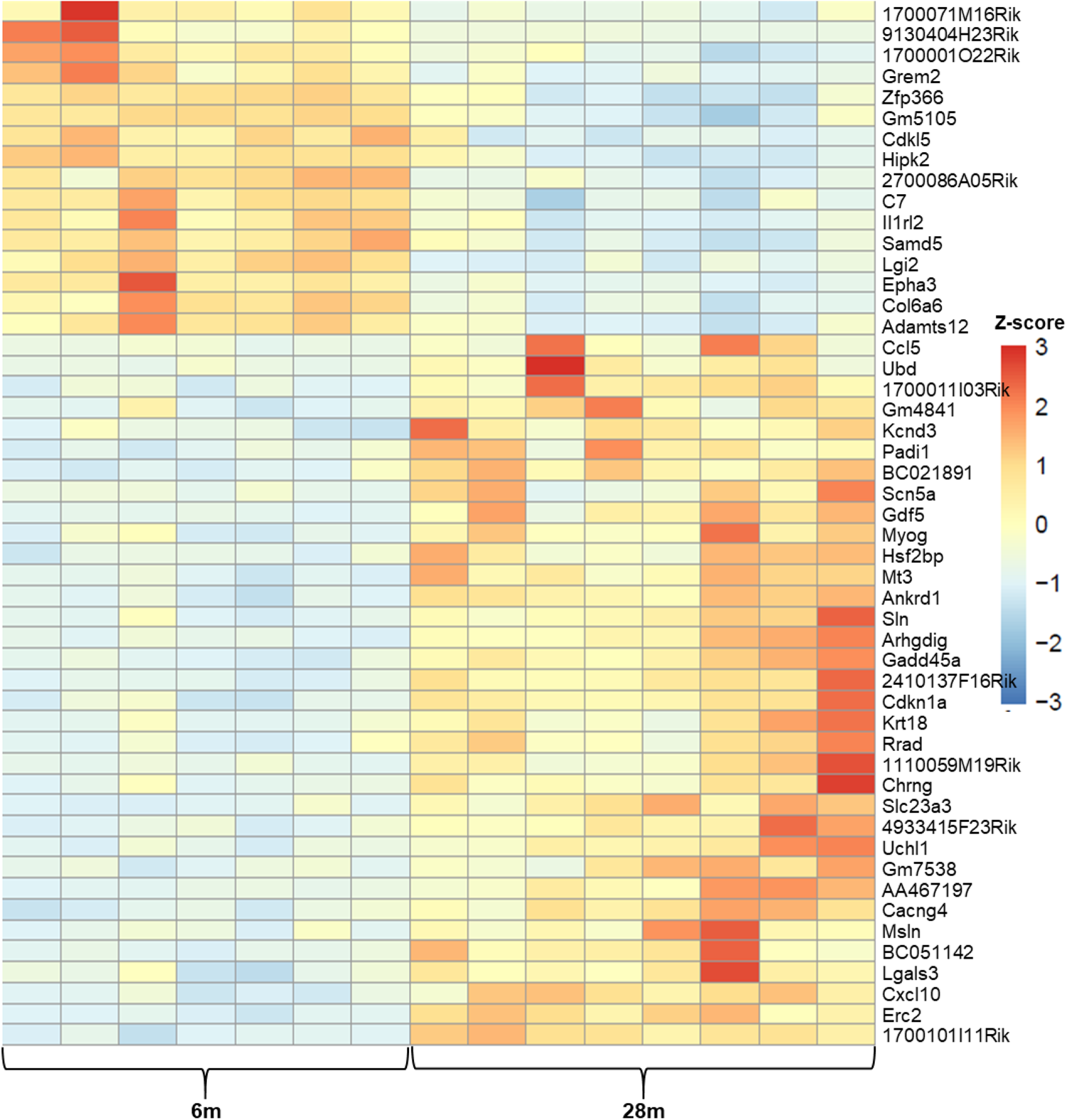
6m vs. 28m Top 50 Z-scores Heatmap. The names of the genes are to the right of each row, and each column = expression data from an individual mouse, 6m = 6 month old and 28m = 28-month old mice, color coded key to fold change z-score in on the right with red the highest (+3) and dark blue the lowest (−3).

**Figure 3.**
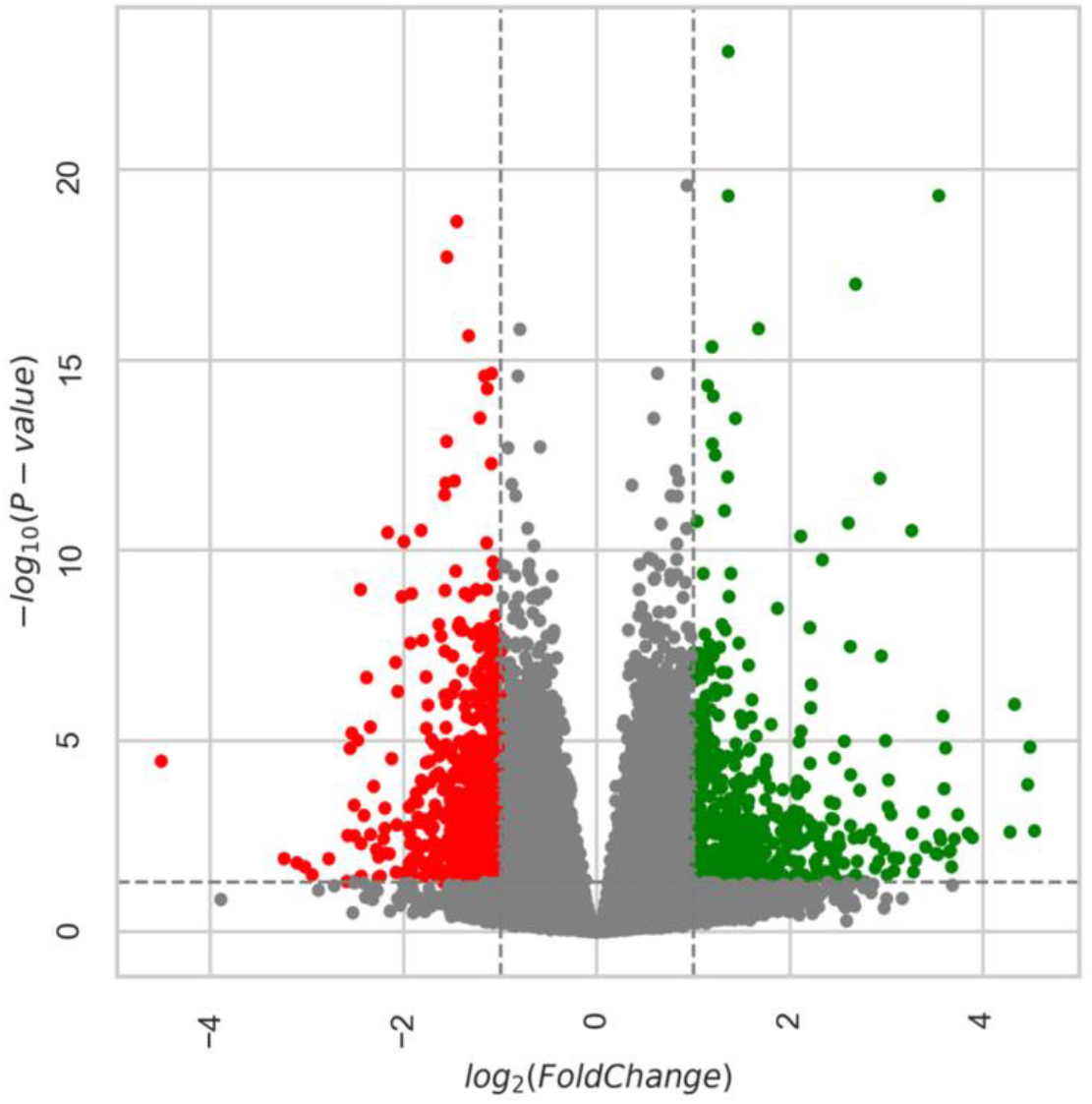
Volcano Plot: 6m vs. 28m. Each dot (red indicates down-regulated gene expression with age and green indicates upregulated) represents one gene with the log2 fold change on the x-axis and the adjusted p-value on the y-axis. Dashed lines indicate the cut-offs of adjusted p-value<0.05 (horizontal line) and log2 fold change >|1| as the two vertical lines. All colored circles were considered significantly different gene expressions with age. 6m = six-month old mice and 28m = 28-month old mice.

**Figure 4.**
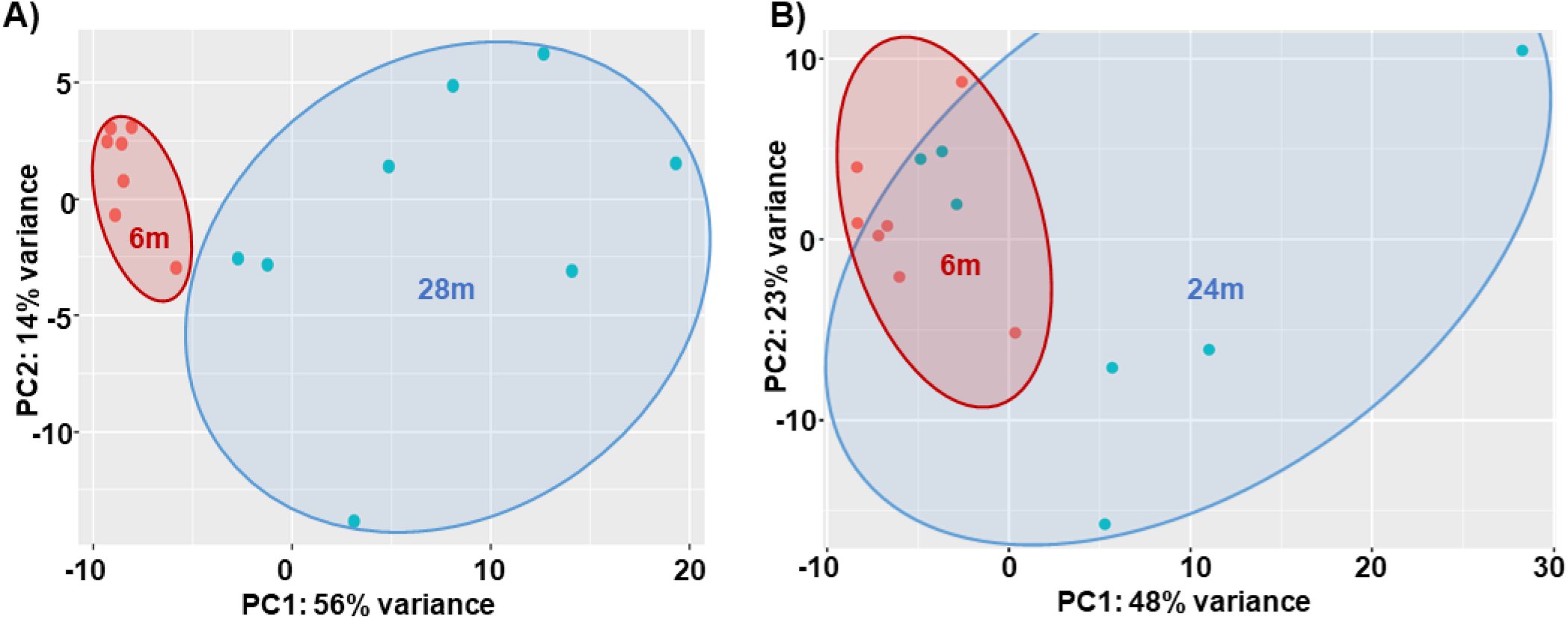
PCA Plots. **A)** 6m vs. 28m, **B)** 6m vs. 24m Key: 6m = 6 months old, 28m = 28 months old, PCA = principal components analysis, PC = principle component, and percent variance indicates how much of the variability between subjects is explained by the components, red dots = 6m and green dots = 28m.

Analyzing the RNAseq data with GSEA, we note that 1049 genes remained of those that fell under the cut-off (|log2fc|≥1, adj. pval<0.05). Using GOrilla to further determine gene set enrichment in this comparison (using the same genes identified in GSEA), there were 73 gene ontology terms enriched (minimum False Discovery Rate q-value, FDR q-val<0.10; 72 terms FDR q-val<0.05) ranging from a high enrichment of 26.29 to a low of 1.12. In all there were 8 gene sets highly enriched, E, (E>10, averaging 17.6±6.0 sd, standard deviation), including: GO:0016907 (G protein-coupled acetylcholine receptor activity, enrichment, E=26.3), GO:0098639 (collagen binding involved in cell-matrix adhesion, E=25.7), GO:0048407 (platelet-derived growth factor binding, E=21.9), GO:0008046 (axon guidance receptor activity, E=14.3), GO:0035373 (chondroitin sulfate proteoglycan binding, E=14.2), GO:0015464 (acetylcholine receptor activity, E=13.6), GO:0005021 (vascular endothelial growth factor-activated receptor activity, E=12.9), and GO:0030020 (extracellular matrix structural constituent conferring tensile strength, E=12.0). There were 359 genes identified by the intersection of GSEA and GoRilla 6m vs. 28m comparison. For further details see **Figure S1, Table S2a and Table S2b** in the Supplement.

#### 24-month-old mice compared to 6-month-old mice

At 24m compared to 6m, there were fewer changes in gene expression than in the 28m (a total of 219 genes changed significantly, adj. p<0.05), with 46 genes decreasing expression and 173 increasing. By expanding to include genes changing with adj. p≤0.05, there were 234 genes, with 5 downregulated (log2fc,≤1; adj. p≤0.05) and 184 upregulated (9 downregulated and 137 with [logfc≥1]; adj. p≤0.05). See **Table S1b** for all genes log2fc≥│1│and adj. p≤0.05. In **Figure 4B**, the 2D principal component analysis (PCA) scores plot indicates an incomplete separation between 24m and 6m clusters, with evident overlap. This result was confirmed by using the supervised multivariate analysis based on a partial least squares-discriminate analysis (PLS-DA) (component 1, 6m, was 23% and component 2, 28m, was 48%).

Analyzing the RNAseq data with GSEA, we note that 127 genes remained of those that fell under the cut-off (|log2fc|≥1, adj. pval<0.05). Using GOrilla to determine gene set enrichment of the same genes identified in GSEA, there were 19 gene ontology terms enriched (FDR q-val<0.10; 18 terms FDR q-val<0.05), ranging from a high enrichment of 43.19 to a low of 1.39. In all there were 3 gene sets highly enriched (E>10, averaging 32.5±11.8 sd), including: GO:0001602 (pancreatic polypeptide receptor activity, E=43.2), GO:0001601 (peptide YY receptor activity, E=34.55), and (neuropeptide Y receptor activity, E=19.81). There were just three genes identified by the intersection of GOrilla and GSEA for this 6m vs. 24m comparison. For further details see **Figure S2, Table S2c**, and **Table S2d** in the Supplement.

### Regressions and Correlations of CFAB and RNAseq

We primarily focused our attention on the changes that occurred in the transcriptome between the adults (6m) and the oldest group (28m). This is because the alterations in the genome were most extreme at the advanced age (more than 6700 genes changed significantly with age), and we know from previous work that the most profound changes in function, muscle health, and contractile ability occur at the older ages in mice (Graber, 2015; Graber, 2013; Graber, 2020). However, we have presented data including linear regressions from the other conditions for full comparison purposes. See the data sets in the Online Only Supplement **Table S4** for more details.

#### 28-month-old mice compared to 6-month-old mice

Regression analysis of the 6m with the 28m determined that there were 689 genes with at least a moderate (R≥0.50) correlation with physical ability (CFAB score), and of these 253 were strongly associated with CFAB (R≥0.70). In **Table 3** we list the top 20 (by R^2^) age-regulated genes associated with physical function. See **Table S4a** for details of all genes with R>0.70 (regression p<0.05), log2fc≥│1│and adj. p≤0.05.

**Table 3.**
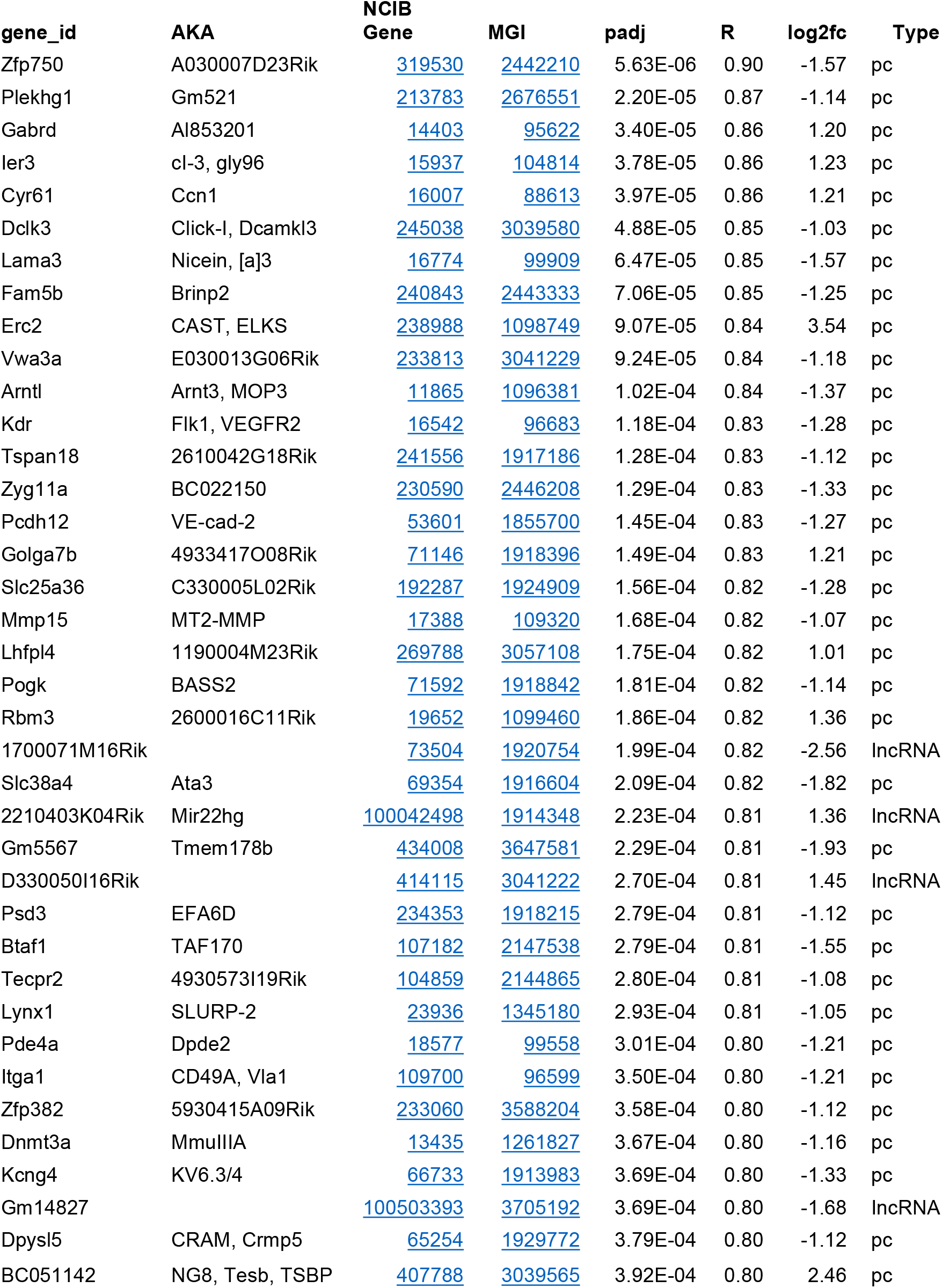
Age-Regulated Genes Associated with Physical Function: 6m vs. 28m (R≥0.80) AKA = also known as, NCIB Gene is from https://www.ncbi.nlm.nih.gov/gene, MGI = Mouse Genome Informatics from http://www.informatics.jax.org/marker, lo2fc = log base 2 fold change, adj. p = multiple comparison adjusted p-value, R = Pearson correlation from simple linear regression.

#### 28-month-old mice vs. 24-month-old mice compared to 6-month-old mice

When we combined the results (genes that changed with age at least log2fc≥1, and adj. p-val.<0.05) from all 3 groups, regression analysis determined that there were 550 genes with at least a moderate (R≥0.50) correlation with physical ability (CFAB score), and of these 108 were strongly associated with CFAB (R≥0.70). See **Table 4b** for details.

**Table 4.** Mouse Characteristics. Body mass is the weight at tissue collection, Total Muscle is the combined mass of the mean extensor digitorum longus, tibialis anterior (TA), gastrocnemius, plantaris and soleus muscle. Statistics are from a simple one-way ANOVA, different letters equal statistical significance at p<0.05 using a Least Significant Differences post hoc test.

| **Age**<br>**months** | **n** | **Body**<br>**Mass**<br>**g** | **Total**<br>**Muscle**<br>**mg** | **TA**<br>**mg** |
| --- | --- | --- | --- | --- |
| **6** | 8 | 33.04±0.42 | 286.75±6.97 | 58.39±1.30 |
| **24** | 8 | 33.71±0.67 | 254.52±7.27<sup>a</sup> | 50.14±1.96<sup>a</sup> |
| **28** | 8 | 31.01±1.05 | 206.13±5.75<sup>b</sup> | 43.48±0.96<sup>b</sup> |

#### 24-month-old mice compared to 6-month-old mice

When examining only the relationship between the results from the 6m and 24m groups, regression analysis determined that there were 55 genes with at least a moderate (R≥0.50) correlation with physical ability (CFAB score), and of these 22 were strongly associated with CFAB (R≥0.70). See **Table S3c** for details.

## Discussion

### Physical Function Declines with Aging

It is well-established that both rodents and humans lose muscle mass and strength as they get older. Alongside this decline in muscle mass and strength comes a decline in physical function and exercise capacity (Graber, 2020; Graber, 2013). Reductions in power production and contractile velocity have been shown to precede loss of strength and mass, indicating that deterioration other than atrophy contributes to the onset of muscle dysfunction (Graber, 2015; Made-Wilkinson, 2015).

Various hypothesis have been proposed to explain the mechanisms of both early onset loss of power and the disconnect between mass retention and strength loss in the context of declining physical performance. Loss of so-called “muscle quality” is one such theory that can be explained via numerous mechanistic avenues. During aging, the infiltration of fat, connective tissue and scar tissue into a muscle can reduce the overall cross-sectional area devoted to contractile units while altering structural parameters of the tissue, and combined with other macro level alterations such as tendon stiffening, may reduce power and strength at the whole muscle level (Wu, 2020; Rahemi, 2015). In addition, at the cellular level, there are many deleterious changes with aging that can affect contractile velocity, power production and force generation, such as: post-translation modifications to key contractile proteins that might inhibit or slow down cross-bridge cycling or limit the number of bound myosin heads at any given moment (such as glycosylation of actin or myosin heavy chain), a shift in the myosin light chains to slower isoforms, fiber-type shift to slower less powerful myofiber types and increased hybrid fibers, enhanced denervation of type 2 motor units with reduced rates of re-innervation, cell signaling abnormalities potentially resulting from such divergent sources as enhanced global inflammation and reduced hormonal signaling, autophagy dysregulation resulting in cellular “junk” accumulation including reduced mitophagy rates leading to a greater number of dysfunction mitochondria leaking enhanced protein/DNA/RNA damaging reactive oxygen species (ROS), and many others (Prochiewicz, 2007; Wiedmer, 2021).

In the current study we set out to obtain a transcriptome profile during aging in the mouse which we could then compare to the decline of physical function with the intention of developing mechanistic hypotheses based upon this relationship. It was clear from the data that there is a vast wealth of correlative connections between various mRNA species relative abundance and the overall state of functional health (measured by CFAB) in our mice. With the vast amount of data, we relied on Gorilla and GSEA analysis of gene ontology to get some clues as to what the genes associated the most with functional decline seemed to be telling us about the cellular process in flux. Calcium handling was one process that seemed to jump out immediately. We also found evidence of denervation, neuromuscular junction dysfunction, and motor neuron alterations. In addition, we suspected a priori that mitochondrial changes would be highly relevant, but the evidence we found for this was less.

### Aging and the Transcriptome

Even though our cut-off point for significant gene expression changes was log2fc≥1 (2-fold), that does not mean that genes below that cut-off do not play a role in age-related functional loss. To narrow down the more than 6500 genes found to change between adult and older mice, we chose a standard two-fold change in gene expression as both real change and clinically significant. In **Table 1** of the supplement we list the top 30 mRNAs that increase with age and in **Table 2** the top 30 that decrease with age (from 6-28 months). Note that some of these are also correlated with declining function, but some are not.

### Transcription Factor Gene Expression with Aging

A recent study narrowly focused on multi-tissue conserved epigenetic regulation and the transcriptome (genes within 5000 base-pairs of transcription start sites) used the quadriceps of 6 and 24-month old mice as part of their analysis (Sleiman, 2020). It is interesting to note that while they only report and investigated a very narrow scope of genes, many of their top genes were not shown to increase or decrease significantly in our TA transcriptome within our cut-offs (adj. p<0.05 and log2fc≥1) at 6 and 24 months—perhaps partly indicating a likely difference between the highly glycolytic fast twitch TA and the more oxidative quadriceps requiring a nuanced approach to this type of analysis due to the heterogeneous nature of individual muscles.

However, if we instead compare 6 month old mice TA to 28 month old mice TA, we do see some evidence of transcriptional regulation alterations similar to Sleiman et al. By widening our scope to significance at adj. p<0.05 and any log2fc, we see similar results for transcription factors related to aging such as the SREBF family motifs (SREB1 log2fc −0.61, and SREB2 log2fc −0.67, both adj. p<0.001) that are known to regulate lipid homeostasis, and Mecp2 (log2fc −0.30, p=0.006) that represses expression of genes and is a regulator of normal neuron function. We also see significant age-associated expression changes of members of the Zbt family (log2fc: Zbtb37 - 1.06, Zbtb7c −0.63, Zbtb46 −0.73, Zbtb22 0.25, Zbtb48 0.31, Zbtb33 −0.29, Zbtb10 −0.33, Zbtb5 −0.30, Zbtb11 −0.22, and Zbt20 0.34), that code for the zinc finger and BTB domain-containing proteins which are known as transcriptional repressors. For example, ZBTB20 promotes production of pro-inflammatory cytokines by downstream activation of Toll-like receptors. In support, we uncovered that Nfkbia (Nuclear factor of kappa light polypeptide gene enhancer in B-cells inhibitor, alpha) increased expression by log2fc 0.96, and thus could be involved int reducing NF-κB transcription factor (confirmed significant reduced expression of Nfkb1 is log2fc −0.45). Therefore, upregulation of ZBTB20 in our older mice would support increased inflammation—one hallmark of aging and one of the likely mechanisms contributing to muscle atrophy.

### Age-Related Gene Expression Relationship with CFAB

Using the single CFAB score to represent generalized physical function at the three different ages we ran linear regression analysis to determine which of the age-associated gene expression changes correlated at a moderate or strong level. It is important to note that there are differences between the linear regressions at the various ages. This is not surprising because some of the genes did not change with age at 24-months but did at 28-months. In addition, gene expression changes that are potentially associated with physical function over the lifespan (6-to 24-to 28-months combined) may not be the same as gene expression changes associated with physical function when comparing only two of the age groups (6 to 24, or 6 to 28 months, for example). Gene expression at one of the ages may not affect function in the same way as at the other. One example would be Erc2 (also known as Cast1), a gene that codes for the protein ELKS-Rab6-interacting protein 2, which has many roles including organization of the cytoskeleton structures involved in pre-synaptic vesicle release (Ko,2003; Chen, 2011). Erc2 has a log2fc 3.41 between 6 and 24 months, and then log2fc 3.54 between 6 and 28-months. Overall, the linear regression of Erc2 with CFAB of all three age groups of mice has a Pearson correlation of R=0.78, with only the 6-month and 24-month R=0.64, and finally with only the 6-month and 28-month R=0.84. Thus, our hypothesis is that levels of Erc2 mRNA become more critically related to physical function at advanced ages. Since Erc2 is involved in the organization/fusion of vesicles at the presynaptic terminal of nerves (presumably vesicles containing acetyl choline in the α-motor neuron in this case), it makes sense that the older the mouse, likely the more critical the role of Erc2 to neuromuscular performance might become.

### Potential Mechanisms of Functional Age

Through examination of the genes most affected by aging (see **Table 1 and 2**), and also those with the highest correlation with CFAB (See **Table 3**), we note some common themes related to the protein function of some of the top gene changes: calcium handling dysregulation (Sln, sarcolipin with log2fc 4.33), denervation (Achg log2fc 3.599) and neuromuscular junction degeneration, and proteolytic process regulation (Ubd, ubiquidin log2fc 4.46). In the next few sections, we will examine the significance of these changes in more detail.

### Denervation and Neuromuscular Junction Degradation

The acetylcholine receptor (AChR) in skeletal muscle receives acetylcholine diffusing across the synaptic cleft after being exported via exocytosis from the motor end plate of the innervating motor neuron when an action potential is propagated (back to that process in a moment). AChR has 5 different subunits: α, β, δ, ε and γ. The complex consists of 2 α, 2 β, 1 δ, and 1 ε (in mature muscle cells) or γ (in embryonic or denervated myofibers). Chng (acetylcholine receptor subunit gamma) is only expressed in mature skeletal muscle after denervation (Ma, 2005)—with the ε subunit returning long after denervation (Adams, 1995). There was a log2fc of 3.599, equivalent to a 1211% increase, in Chng in 28-month old mice compared to 6-month old mice (R=0.46 with CFAB); but Chne (epsilon subunit) was increased only log2fc 0.50, indicating an 860% relative increase of Chng versus Chne that suggests increased flux of denervation (See **Table S4** for all Chng and associated gene expression) in the older animals. Denervated muscle fibers in older animals trend to be less robustly reinnervated than in younger animals, resulting in eventual myofiber death and muscle atrophy (Hepple, 2016) Additionally, in older animals a fiber type shift to a more type 1 slow twitch fiber composition occurs as former type 2 muscle fibers that are denervated are more often reinnervated with type 1 motor neurons. This combination of switching to less powerful myofiber type, coupled with an overall loss in the total number of fibers (not to mention atrophy from other causes) may lead to a reduction in peak power generation in older muscle (Graber 2015).

Formation of the motor endplate, in particular the clustering of ACHRs is propagated by the release of agrin by a motor neuron, that binds to the MuSK receptor (and dystroclycan and laminin to form a stabile scaffold), causing MuSK to phosphorylate and to downstream activate and recruit casein kinase 2 (Csnk2), rapsyn (Rapsn), and Dok-7 to form the ACHR clusters. Motor neuron outgrowth and attachment to myofibers is dependent upon the expression of MuSK and Agrin at the motor end plate (Dimitropoulou, 2005). Musk and Agrin genes are both downregulated significantly in older mice (log2fc −0.627 and −0.390, respectively). Agrin acts to stabilize the MuSK receptor to the extracellular matrix and the cytoskeleton forming a focal point for ACHR clustering (Swenarchuk, 2019). Additionally, DOK4 is a peptide involved in neuronal outgrowth (gene log2fc −0.64) upstream of Rap1 (a g-coupled protein) and the ERK pathway. Interestingly, Trim9 (log2fc −2.51) is a negative regulator of synaptic vesicle transmission that acts as a ubiquitin ligase to regulate the SNARE complex formation, and is important for axon guidance.(Berti, 2002; Plooster, 2017)

### Calcium Handling Dysregulation (Implications and Effects)

Increased calcium levels in the sarcoplasm can significantly alter numerous signaling pathways and mechanisms that rely upon Ca^+2^ as a second messenger. Most obviously the primary example in skeletal muscle is promotion of contraction induced by the binding of Ca^+2^ to troponinC, which then causes a conformation shift of the troponin complex that moves tropomyosin away from the myosin binding site on actin, allowing for cross-bridge cycling. Normally this process is induced by a calcium influx from the sarcoplasmic reticulum when the ryanodyne (RYR) receptor is prompted to open by the voltage-gated dihydropyridine receptor responding to a propagating action potential. Relaxing of the contractile elements is induced when Ca^+2^ disassociates from troponinC as the sarcoplasmic endoplasmic reticulum ATPase (SERCA) pumps Ca^+2^ against the concentration gradient from the sarcoplasm back into the sarcoplasmic reticulum using ATP.

Sarcolipin (Sln, increased log2fc 4.33, adj. p=1.1×10^−6^) is one of the regulatory elements of SERCA, along with phospholamban (Pln, increased log2fc 0.663, adj. p=0.0003), and myoregulin (Mrn, aka 2310015B20Rik, did not significantly alter), that functions by blocking the pumping ability of SERCA even while allowing ATPase activity to consume energy and generate heat as a byproduct (Anderson, 2015). SLN and PLN are additive in effect and can cause super-inhibition of SERCA pumping activity when expressed in the same cell. Additionally, there are other midcropeptides newly identified that are involved in muscle regulation of SERCA including DWORF (increasing SERCA pumping) and the negative regulators endoregulin and another-regulin (Anderson, 2017). This increase in expression may have implication as an adaptive strategy for increasing non-shivering thermogenesis to ward off body temperature dysregulation in older mammals and/or to improve energy balance in more sedentary individuals (Bal, 2012); but may well have adverse consequences concerning muscle and physical function. SLN expression is not only greatly over-expressed in 28-month old mice (log2fc=4.33) but is negatively correlated (R=−0.55) with CFAB functional scores. One mechanism by which this could occur is by increasing the time needed to relax muscle fibers between contractions, by delaying disassociation of Ca^+2^ from troponin due to an increased sarcoplasmic calcium concentration, which would potentially lead a decrease in power production. Sarcolipin is overexpressed in Duchenne muscular dystrophy (DMD) patients and DMD transgenic mouse models, and the knockdown of SLN restores muscle and physical function (Voit, 2017). However, knock-out of SLN prevents normal hypertrophic and fiber-type shift response to overloading, and increases ½ relaxation rate compared to wild type (Tupling, 2011; Fajardo, 2017). Transgenic mice over-expressing SLN have been shown to have an increased metabolic rate, ½ relaxation time, while increasing SLN in rat muscle has been shown to decrease both maximal isometric force and ½ relaxation time, bolstering this theory (Maurya, 2015; Tupling, 2011, Tupling, 2002).

In addition to dysregulating cross-bridge cycling and force generation, overexpression of SLN and PLN leading to increased prevalence of cytosolic Ca^+2^ abundance may stimulate numerous calcium-dependent signaling pathways. For example, increased levels of sarcoplasmic calcium can decrease promoter activity for CGRP (calcitonin gene-related peptide), which is alternatively spliced from the calcitonin gene. CGRP binds to the calcitonin receptor like receptor (CKACRL) which consists of three different subunits: the receptor component protein (Rcp), the calcitonin like receptor (Calcl, log2fc−0.34, adj. p=0.006), and the receptor activity-modifying protein 1 (Ramp1). Ramp1 (log2fc 0.53, adj. p=0.01) is involved in angiogenesis and wound healing.

### Disuse Atrophy

As reported by Mahmassani et al. 2019, after a 5-day period of bedrest 61 genes were differentially expressed (pre-post) in the vastus lateralis of younger adults compared to older, with 51 of these genes changing only in young adults to levels equivalent to older adults at baseline, suggesting that in some ways that older muscle resembles adult muscle suffering from disuse atrophy. In our study we determined that of the top 10 genes they touted as being differentially upregulated in younger mice during bedrest, in our oldest mice Fasn, Pfkfb3, and Rps4x were significantly expressed differentially from adult mice with log2fc of −0.679, −1.09, and 0.93, respectively. However, in their top 10 downregulated genes in adult humans after bedrest, only Nov (−0.545 log2fc, trend adj p-val=0.067), Apln (0.81 log2fc, trend adj p-val=0.067), and Myl12a (1.39 log2fc, trend adj p-val=4 ×10^−10^) were significantly altered in our 28 month old group compared to the 6-month. Fisher and colleagues (Fisher 2017) used tetrodotoxin administration as a model of reversible denervation-induced disuse atrophy and demonstrated that there was a time course dependent relationship for various gene expression changes with four of their top 7 differentially expressed atrophy-related genes also showing significant changes in our 28+month old mice, further making the claim that older muscle dysfunction may partly be due to chronic disuse patterns. This presents an intriguing concept for future deliberation to determine which elements of acute detraining/disuse could be contributing to long-term disuse atrophy in older adults and which of these might be ablated by minimal increases in activity rates or other interventions to preserve function.

### Caveats

First of all, it is well-established that alterations in gene expression are often not equivalent to alterations in protein expression: in effect, the transcriptome ≠ proteome! Thus, it will be important to investigate protein abundance of physiologically relevant gene expression changes to determine the true extent of influence any of these proposed mechanistic components contributing to declining function, and, furthermore, to establish cellular signaling mechanisms connecting the numerous potential sequences of events. In addition, this study has a relatively small n, which makes correlation and linear regression association less reliable. This study only included male mice, so it will be necessary for future work to investigate whether there are any sexual dimorphisms in aging gene expression patterns.

Another limitation of the current study is that we only have three age groups to draw conclusions from, and thus having less accuracy in determining changes over the lifespan. Determining which changes are early onset will require middle-aged groups (16-20 months), and adding an oldest-old (e.g. 32+-month old group) would allow us to gain insight into potential mechanisms related to successful aging of the oldest-old.

Despite these limitations we believe this data set is novel and comprehensive. We have performed transcriptomics at three different age time points. Furthermore, we have performed a comprehensive battery of physical function tests at each time point. The combination of transcriptome data with functional data is novel and will be valuable in establishing the framework and preliminary data to begin designing mechanistic studies of key genes of interest.

### Future Directions

Establishing physiological relevance with concurrent changes in protein expression is important and will help us design mechanistic studies to establish cause and effect for genes/proteins that induce sarcopenia. We will also seek to determine how interventions can alter the transcriptome to potentially create a “younger” transcription profile than would be expected by biological age, and whether this would translate into improvements in functional capacity. Exercise, in its many forms, is one intervention known to improve function, and comparing the transcriptome of exercise and control mice over the lifespan would be a valuable way to assess which genes important for maintaining functional are modulated via exercise, and, conversely, which genes do not change expression from an exercise treatment.

With the ubiquitous use of the mouse model in aging, mechanistic, and pharmaceutical research, understanding both parallels and differences in age-associated gene expression with humans is a necessary future undertaking. A recent comparative study of gene array data of skeletal muscle in mice and humans revealed 249 homologous overlapping age-related genes (Zhuang, 2019), but noted 6333 differentially expressed skeletal muscle genes between under 30 year old and over 65 year old humans—very similar to our finding of 6587 in 6-month to 28-month old mice. It is important to note that, as we have uncovered in this study, the age of the older mice plays a key role in differential gene expression. According to our data, mice experience a rapid transcriptomic change between 24 and 28 months of age suggesting that mice at the older age are experiencing far more age-related changes than younger mice. In the Zhuang and colleagues study the mice ages from the gene arrays they investigated were not given. Thus, more research is needed to establish age-associated gene expression changes related to functional decline in humans and which of these overlap with mice.

## Conclusion

This current study is a first step in investigating potential novel mechanisms of age-related functional decline manifested in differential gene expression with aging. More work is needed to determine the physiological relevance of the many changes uncovered and to determine any proteomic alterations predicated by the altered gene expression. The data sets we present herein will help to identify and characterize cellular mechanisms responsible for how age induces a decline in muscle health and physical function with the potential for uncovering novel therapeutic targets.

## Experimental Procedures

### Mice

Three different ages of C57BL/6 male mice were obtained from the National Institutes of Health National Institute on Aging Charles River Aging Rodent Colony (a subset of mice from the previously published Graber, et al. 2020 were randomly selected for this study: n=8 for all at 6-months-old, 24-months-old, and 28+-months-old). The characteristics of the mice are presented in **Table 4**. Mice were treated humanely under approved IACUC protocols and were group-housed at 22 °C with a 12-hour:12-hour light/dark cycle. Food and water was provided ad libitum.

### Functional Testing

We used the protocols described in prior work (Graber, 2020) to measure the physical function and exercise capacity of the mice using our CFAB composite scoring system. In brief, CFAB defines function using a composite of 5 well-validated functional tests: rotarod for overall motor function (Graber, 2013), inverted cling for four-limb strength/endurance (Graber, 2013), voluntary wheel running as a measure of volition exercise and activity rate (Graber, 2015), grip test to measure fore-limb strength (Graber, 2018), and treadmill running for aerobic endurance (Graber, 2019). Using adult 6-month-old mice mean as the control reference, the distance in units of standard deviation (SD, calculated from 6-month-old group) of the score of each individual mouse for each test from the adult mean (standardized score) was calculated. All five standardized scores of each individual mouse are then summed to produce the CFAB score of that mouse. A further brief discussion of the functional measurement in the **Online Supplemental Procedures Section**.

### Tissue Collection and Handling

At the completion of the testing protocols the mice were euthanized after non-survival surgery to collect the hindlimb muscles. The muscles were blotted dry, weighed, and then immediately flash frozen in liquid nitrogen. Subsequently the muscles were stored at −80 °C until total RNA extraction.

Total RNA extraction has been previously described (Graber, 2017). In brief, we used Tri-Reagent (Molecular Research, #TR118) using the manufacturer’s instructions to extract total RNA from TA muscle, using the entire TA muscle. We quantified the extraction using a Nanodrop2000 (ThermoScientific), with mean concentration 330.8 ± 24.3 ng/µl, 260/280 ratio 1.69 ± 0.020, 260/230 ratio 1.98 ± 0.09. RNA integrity was determined using an Agilent Bioanalyzer 2100, mean RIN was 9.23 ± 0.139. Two isolated RNA samples of the 24 total (n=1 each from 6m and 24m groups) did not meet the standard lower limits for purity and integrity and were not used for RNAseq.

### NGS RNAseq

RNA samples (n=22 total; n=7 6m, n=7 24m, and n=8 28m) were quantified using a Qubit fluorometer and qualities were assessed with an Agilent Bioanalyzer. Poly-A+ RNA was enriched from ∼0.5 ug of total RNA and used as a template to generate sequencing libraries using the New England Biolab NEBNext Ultra RNA Library Prep Kit following the supplier’s protocol. Libraries were pooled and sequenced on an Illumina NextSeq 550 High-output flow cell with the 75 base pair single-end protocol. Raw NGS data is stored at The Geo record GSE152133.

### Data Analysis

#### General

We used SPSS v24 and v25 (IBM) to analyze the statistics. Data reported as means ± standard error, unless otherwise designated. Significance was designated as p<0.05. Linear regression and Pearson’s Correlation were used to establish relationships between gene expression and CFAB. Depending upon the comparison we used either ANOVA, or ANCOVA (adjusting for body mass). Post-hoc analysis for ANOVA and ANCOVA used least significant differences. RNAseq data analysis is outlined below.

#### RNAseq

The reads in fastq format were aligned to the mouse mm10 genome using the splicing aware software STAR, version 2.5.4b, using the ENCODE recommended parameters. The genome index was built with the Illumina iGenomes UCSC mm10 genomic sequence and annotation file, and reads mapped to genes were quantified with the STAR –quantMode GeneCounts option. The read counts per gene for each sample were input into the DESeq2 differential expression program. Following the DESeq2 vignette, differentially expressed genes were called with a adjusted p-value cut-off of less than 0.05 and a log2 fold-change of +/-1.0. The rlog function in DESeq2 was used to generate a table of log2 normalized counts, which was used to generate the PCA plots and heatmaps. The heatmap program was used to create the heatmap figures. The principle components analysis determined the gene sets that contributed most to the variability between the different aged groups and for which genes contributed most to explaining CFAB variation. See **Figure 4**.

### Further Data Analysis of RNAseq data and CFAB Data

The Bioinformatics and Analytics Research Collaborative (BARC) at the University of North Carolina at Chapel Hill performed the following data analysis as consultants to the project: GSEA (Gene Set Enrichment Analysis) was conducted using R referring to the method explained in https://stephenturner.github.io/deseq-to-fgsea/ against NGS datasets (Subramanian, 2005). The reference database used was ‘MousePath_GO_gmt.gmt’ downloaded from http://ge-lab.org/gskb/. Based on the results of GSEA, genes from the NGS datasets with the cutoff (|log2fc|≥1, adj. pval<0.05) were further filtered. 127 and 1049 genes were left from adult vs. older comparison and adult vs. elderly comparison respectively.

Then, GOrilla analysis was rendered on http://cbl-gorilla.cs.technion.ac.il/ against the same two datasets as used by GSEA. Enrichment is the over or under representation of differentially expressed genes in functional categories (the GOs/ gene ontologies). GOs can be thought of as bins of genes that are part of a pathway. In GOrilla, a statistically significant enrichment score (all are positive) indicating this pathway is activated (Eden, 2009).

The final step was to intercept the results from both GSEA and GOrilla. The intersection simply provides high confidence between two approaches for ascribing functional categories to the data. GSEA and GOrilla try to do similar things but have different methods. GOrilla is the older tool and more traditionally used and focuses on the significant genes, whereas GSEA considers all of the genes in an experiment, not only those above an arbitrary cutoff in terms of fold-change or significance. Moreover, GSEA assesses the significance by permuting the class labels, which preserves gene-gene correlations and, thus, provides a more accurate null model.

## Acknowledgements

The authors would like to acknowledge the excellent work of our statistical consultants at the University of North Carolina-Chapel Hill School of Medicine Bioinformatics and Analytics Research Collaborative (BARC) for providing technical support, and especially Corbin D. Jones and Paul Grant. We also want to acknowledge the valuable contribution of Nainika Nandigama, and Alyssa Fennel who helped compile some spreadsheets for the Online Supplemental Material.

## Author Contributions

Conceptualization: T.G.G., B.B.R.; methodology: T.G.G., J.T., and S.W.; validation: T.G.G., formal analysis: T.G.G., Z.M, M.L.P. J.T., and S.W.; investigation: T.G.G., R.M., J.T., and S.W.; resources: T.G.G., and B.B.R.; writing—original draft: T.G.G.; writing—review and editing: T.G.G., B.B.R., R.M., J.T., S.W., Z.M., M.L.P.; supervision: T.G.G.; project administration: T.G.G.; funding acquisition: T.G.G.

## Data Sharing

The raw NGS RNAseq data is deposited in GEO (GSE152133). We have provided differential expression spreadsheets in the Supplementary Files available on Figshare at **PUBLIC DOI PENDING**. Other data will be provided upon reasonable request to the corresponding author.

## Supporting Information Listing Section for

Age-Related Transcriptome Alterations in Skeletal Muscle Related to Physical Function Decline of Older Mice

## Table of Contents

***Full NGS RNAseq Raw data at GEO GSE152133***.

Supplementary Information available via Figshare at: PUBLIC DOI PENDING

### Supplementary Figures (PDFs)

**Figure S1** GoRilla Gene Enrichment: 6m vs. 28m

**Figure S2** GoRilla Gene Enrichment: 6m vs. 24m

### Supplementary Tables (Excel Spreadsheets)

**Table S1** Gene Expression Changes with Age

**Table S2** GSEA and GOrilla GO Enrichment

**Table S3** GSEA and GOrilla_gene set intersection

**Table S4** Linear Regressions of Age-Changing Genes with CFAB

### Supplementary Text

Online Supplemental Procedures Section (PDF)

