## Supplemental Methods for "Skeletal Muscle Transcriptome Alterations Related to Physical Function Decline in Older Mice"

**Online Supplemental Procedures Section**

**CFAB** (**C**omprehensive **F**unctional **A**ssessment **B**attery). CFAB is a composite scoring system consisting of 5 different well-validate non-colinear functional tests (activity level/volitional exercise rate via voluntary running wheels, rotarod, grip testing, inverted cling grip test,, treadmill endurance test) (Graber, 2020; Graber, 2019; Graber; 2018; Graber, 2015; Graber, 2013). The system uses the average 6-month old adult C57BL/6 mice performance in each test as the baseline, and measures how far an individual mouse's performance is from that baseline in units of standard deviation (derived from the 6-month old mouse data set). The deviation in performance for each of the 5 tests in sd is summed to produce the CFAB score.

Each individual functional test was performed at approximately the same relative time daily for each mouse in order to maintain equality in circadian rhythm disruption (after 3pm).

**Functional Tests Comprising CFAB:**

A) *Activity/Volitional Exercise Rate* (voluntary wheel running): To track activity, the mice were singly housed with a running wheel (Columbus Instruments) for 1 week. The number of revolutions of the wheel over the week were converted to km/day and reported as such. NOTE: because of the relatively small diameter of these wheels which are designed for barrier cages and fit inside the homes cage, the mice do not generally run as far per day as some other wheel systems (Graber, 2020)

B) *Rotarod:* To quantify overall neuromuscular function (endurance, power production, balance and coordination) we used a Panlab Rota-Rod. The procedure involved two training sessions, one per day (three trials per day, minimum of 15 minutes rest between trials), to acclimate the mice to the device, followed by a testing day in which three trials were performed, once again with minimum of 15-minute rest periods between trials. The maximum number of seconds the mouse remained on the rotarod before falling was the outcome measure.

C) *Grip Test Meter:* To directly measure grip strength of the mice, we used a Bioseb grip strength meter. Five trials for forelimb grip was performed in one session. For one trial, the mouse was removed from its cage, held by the tail and placed, gently, so that its paws can grab the bar/grid. Then the mouse was smoothly pulled backed until it releases the bar/grid. We report the highest of the five trials in Newtons as the outcome measure.

D) *Inverted Cling Grip Test:* This test was used to quantify muscle strength and endurance. The mouse gripped a grid (a custom-built device is used) and was inverted. The outcome is latency until falling, in seconds, to the padded surface below, best out of two trials (7 minute ceiling). The mice were tested two times on one day with a minimum of 15 minutes between trials for a rest period. If a mouse held on for less than 10 seconds it was be immediately retested to determine if the fall was a slip. If so, that trial did not count towards the two maximum trials

E) *Treadmill:* The mice were tested for endurance) by running on a treadmill (Columbus Instruments 6 mouse treadmill). The outcome measurement is the length of time run in seconds.

The mice were introduced to the device gradually with two acclimation trials over 2 days. Initially in the first session the mice were introduced to the device and learned to walk, then in the second session the first session was repeated, followed by a second session where the speed was accelerated as they learn to run. During actual testing (day 3) the mouse starts at 3 m/minute and the treadmill is accelerated at 6cm/sec/20seconds until the mice reached exhaustion/failure, with the outcome measure as the number of seconds before failure.
